# Development and validation of an LC-MS/MS method for determination of B vitamins and some its derivatives in whole blood

**DOI:** 10.1101/2021.01.18.427110

**Authors:** David Kahoun, Pavla Fojtíková, František Vácha, Eva Nováková, Václav Hypša

**Affiliations:** Institute of Chemistry, Faculty of Science, University of South Bohemia in České Budějovice, Branišovská 1760, 37005 České Budějovice, Czech Republic; Department of Parasitology, Faculty of Science, University of South Bohemia in České Budějovice, Branišovská 1760, 37005 České Budějovice, Czech Republic

**Author notes:** **Corresponding author:** Full postal address: David Kahoun, Institute of Chemistry, Faculty of Science, University of South Bohemia in České Budějovice, Branišovská 1760, 37005 České Budějovice, Czech Republic, E-mail address.

**Keywords:** B vitamin, Whole blood, Protein precipitation, Liquid Chromatography, Mass Spectrometry

## Abstract

Obligate symbiotic bacteria associated with the insects feeding exclusively on vertebrate blood are supposed to complement B vitamins presumably lacking in their diet, vertebrate blood. Recent genomic analyses revealed considerable differences in biosynthetic capacities across different symbionts, indicating that levels of B vitamins vary across different vertebrate hosts. However, a rigorous determination of B vitamins content in blood of various vertebrates has not yet been approached. A reliable analytical method focused on B vitamin complex in blood and hemolymph can provide valuable informative background and understanding of general principles of insect symbiosis.

In this work chromatographic separation of a mixture of eight B vitamins (B1 – thiamine, B2 – riboflavin, B3 – niacin, B5 – pantothenic acid, B6 – pyridoxine, B7 – biotin, B9 – folic acid and B12 – cyanocobalamine), four B vitamin derivatives (B3 – niacinamide, B6 – pyridoxal-5-phosphate, B6 – 4-pyridoxic acid and B9 – tetrahydrofolic acid) and 3 stable isotope labelled internal standards (B3 – niacin-13C6, B5 – pantothenic acid (di-β-alanine-13C6,15N2) and B7 – biotin-(ring-6,6-d2)) on C30 column was developed. Detection was carried out using dual-pressure linear ion trap mass spectrometer in FullScan MS/MS and SIM mode. Matching internal standards with analytes was done according to the results of linearity, accuracy and precision. Except for vitamin B9 (tetrahydrofolic acid) instrument quantitation limits of all analytes were ranging from 0.42 to 5.0 μg/L, correlation coefficients from 0.9997 to 1.0000 and QC coefficients from 0.53 to 3.2 %.

Optimization of whole blood sample preparation step was focused especially on evaluation of two types of protein-precipitation agents: trichloroacetic acid and zinc sulphate in methanol. Samples of whole blood prepared in six independent replicates were spiked at 10 μg/L and 100 μg/L level. The best results were obtained for zinc sulphate in methanol, but only nine analytes (B1 – thiamine, B2 – riboflavin, B3 – niacin, B3 – niacinamide, B5 – pantothenic acid, B6 – pyridoxine, B6 – 4-pyridoxic acid, B7 – biotin and B12 – cyanocobalamine) were successfully validated. Accuracy of the procedure using this protein-precipitating agent was ranging from 89 to 120 %, precision from 0.5 to 13 % and process efficiency from 65 to 108 %.

**Highlights:**

- LC-MS/MS method for quantitation of eight B vitamins and four B vitamin derivatives was developed.
- Deproteinization agents trichloroacetic acid and ZnSO_4_/methanol were tested for protein precipitation of whole blood.
- Accuracy, precision and process efficiency were evaluated.
- Successful method validation for seven B vitamins and two B vitamin derivatives in whole blood.

## 1. Introduction

B vitamins are supposed to play a central role in the origin and evolution of obligate symbiosis between insects and bacteria. It has been generally accepted that the main role of the maternally inherited obligate symbionts, so called primary symbionts (P-symbionts), is to provide their hosts with compounds missing in their diets, mainly amino acids, vitamins and cofactors [1]. Two ecological groups of insects are recognized as typical hosts of P-symbionts, namely phytophages (e.g. aphids with *Buchnera)* and hematophages (e.g. tsetse with *Wigglesworthia).* The early experiments on the *tsetse-Wigglesworthia* system showed that elimination of symbionts by antibiotics causes loss of fertility, which can be restored by dietary supplementation with B vitamins [2]. Similar results were reported for human louse Pediculus humanus and its bacterial symbionts [3]. Based on these experiments, it was hypothesized that biosynthesis of B vitamins is one of the main roles of these P-symbionts. The molecular era then brought proofs of genome capacity for B vitamin synthesis in *Wigglesworthia glossinidia* and also some other P-symbionts in blood feeding insects [4, 5, 6]. The lack of B vitamins in vertebrate blood and their provision by P-symbionts, thus became a generally accepted reason for the symbiosis between blood feeding insects and bacteria.

Recently, genomic data became available for several symbionts of hematophagous host. Their overview indicates that the biosynthetic capacities vary across different insects-bacteria associations and different vitamins. For example, biotin seems to be mostly provided by the symbiotic bacteria and several instances of HGT of complete biotin operon into the symbionts suggests crucial role of this pathway in the host-symbiont interaction [5, 7]. On the other hand, many symbionts in blood-feeding insects lack the capacity to produce thiamine and they may even scavenge this compound from the host [5, 8].

Considering these long-lasting debates and the significance of blood feeding insects, it comes as a surprise that very little is known about how the amounts of particular B vitamins vary in blood of different vertebrate species. This example illustrates an importance of a precise analytical assay of B vitamins in blood when addressing particular biological questions.

Vitamins of the B complex (B1 – thiamine, B2 – riboflavin, B3 – niacin, B5 – pantothenic acid, B6 – pyridoxine, B7 – biotin, B9 – folic acid and B12 – cyanocobalamine) are a set of low-molecular-weight water-soluble substances that play a key role in many metabolic pathways. Vertebrates generally maintain a low concentration of B vitamins in the blood, do not store them and are dependent on their dietary content.

Some of the B vitamins can be found in blood or tissues in different forms, vitamers or provitamins. Vitamin B1 may be present as the thiamine monophosphate, diphosphate or triphosphate esters, vitamin B3 as the niacinamide, vitamin B6 as the aldehyde pyridoxal, amine pyridoxamine, their corresponding 5’-phosphate derivatives and the pyridoxic acid as a degradation product, vitamin B9 as the tetrahydrofolic acid and vitamin B12 as methyl-, hydroxo-or adenosyl-derivate of cobalamin.

Analysis of a content of individual B vitamins is widely used in a clinical practice and physiology research. However, each vitamin of the B complex is usually analysed separately (e.g. see [9]) or simultaneously only those of the particular interest [10], according to the individual study needs. Various methods of B vitamins analysis are generally used, including microbiological assays, radio immunoassay, protein binding, spectrophotometry, fluorimetry, chemiluminescence, capillary electrophoresis or highperformance liquid chromatography (HPLC) [11, 12, 13]. Howewer, using different analytical methods for different B vitamins introduces many obstacles into the comprehensive surveys where comparing precise levels is crucial.

Recently, liquid chromatography has been established as the most frequently used emerging method for the determination of B vitamins, with the advantage of multi-analytes detection of most of the water-soluble vitamins in single run. The tandem mass spectrometry (MS/MS) detection shows the best results in sensitivity and specificity. Several methods for simultaneous detection of B vitamins using LC-MS/MS are reported in literature for the analysis of various food matrices, human milk or plasma [14, 15, 16, 17].

Unfortunately, a comprehensive method of the analysis of all B vitamins of the vertebrate whole blood samples is missing. With a respect to the whole blood as a nutrient source for blood sucking arthropods and the fact that about 80 to 90 % of the vitamin B1 is present in red blood cells [18, 19, 20], an analytical method for all B vitamins in the whole blood samples is necessary.

In this work, we present a comprehensive analytical method for separation and detection of B vitamins from a whole blood sample in a single run based on HPLC-MS/MS.

## 2. Materials and methods

### 2.1 Chemicals and reagents

All standards and internal standards (IS) were purchased from Sigma-Aldrich: thiamine hydrochloride (CAS: 67-03-8, purity: CRM, abr.: B1), riboflavin (CAS: 83-88-5, purity: CRM, abr.: B2), niacin (CAS: 59-67-6, purity: CRM, abr.: B3), niacinamide (CAS: 98-92-0, abr.: purity: CRM, B3-AM) calcium-d-pantothenate (CAS: 137-08-6, purity: CRM, abr.: B5), pyridoxine hydrochloride (CAS: 58-56-0, purity: CRM, abr.: B6) 4-pyridoxic acid (CAS: 82-82-6, purity: ≥98%, abr.: B6-4PA) pyridoxal 5’-phosphate hydrate (CAS: 853645-22-4, purity: ≥98%, abr.: B6-5P), biotin (CAS: 58-85-5, purity: CRM, abr.: B7), folic acid (CAS: 59-30-3, purity: CRM, abr.: B9), tetrahydrofolic acid (CAS: 135-16-0, purity: ≥65%, abr. B9-THF), cyanocobalamine (CAS: 68-19-9, purity: CRM, abr.: B12), nicotinic acid-^13^C_6_ (CAS: 1189954-79-7, 100 μg/mL in methanol, abr. IS-B3), calcium pantothenate-^13^C_6_, ^15^N_2_ (di-β-alanine-^13^C_6_, ^15^N_2_) (CAS: N.A., purity ≥97%, abr.: IS-B5) and biotin-(ring-6,6-d2) (CAS: 1217481-41-8, purity ≥97%, abr.: IS-B7). Compounds were stored in original packages at −75°C prior to use.

Acetonitrile (LiChrosolv, hypergrade for LC-MS), methanol (LiChrosolv, hypergrade for LC-MS), formic acid (eluent additive for LC-MS), ammonium formate (eluent additive for LC-MS), zinc sulfate heptahydrate (ACS reag. ≥99%) and trichloroacetic acid (ACS reag. ≥99.5%) were purchased also from Sigma-Aldrich. Deionized water was prepared using a purification system Thermo Scientific Smart2Pure 6 UV/UF.

### 2.2 Standard, internal standard and calibration solutions

Standard stock solutions were prepared individually by dissolving appropriate amount of each compound in methanol (B1, B2, B3, B3-AM, B6 and B12), in 1 % HCOOH (B5), in 0.5% NH4OH (B7), in 4% NH4OH (B9), in 0.01M HCl (B6-4PA), in 50% methanol (B6-5P) or in water (B9-THF) to obtain concentration approx. 1000 mg/L (except for B2, B6-4PA, B6-5p and B9-THF where the concentration was 10 mg/L). All standard stock solutions were stored at −75°C. The standard mixture solution A (10 mg/L) was prepared daily in a glass volumetric flask (5 mL) by diluting the calculated amount of each standard stock solution with water and then diluted to obtain the standard mixture working solution B (1 mg/L).

Internal standard stock solutions were prepared individually by dissolving appropriate amount of each compound in 1 % HCOOH (IS-B5) or in 0.5% NH4OH (IS-B7) to obtain concentration approx. 200-500 mg/L. Stock solution of internal standard IS-B3 was obtained commercially as an ampule of 1 mL solution (100 mg/L in methanol). All internal standard stock solutions were stored at −75°C. The internal standard working mixture solution was prepared daily by diluting of 100 μl of each internal standard stock solution with 4.7 mL of water to obtain concentration 2-5 mg/L.

Calibration solutions were prepared daily in brown glass HPLC crimp vials (1.8 mL) diluting the standard working solution B (5 – 500 μL) and the internal standard working solution (100 μL) with water (400 – 895 μL) to obtain 9 different concentrations: 1; 2.5; 5; 10; 25; 50; 100; 250 and 500 μg/L. Total volume of each calibration solution was 1000 μL. Calibration solutions were analysed in triplicate.

### 2.3 Sample collection and preparation

Whole blood samples used for the method development and validation were collected by the licensed practical nurse into 3 mL blood collection tubes containing an anticoagulant potassium ethylenediaminetetra-acetic acid (K3EDTA) purchased from Dialab. The blood collection tubes were mixed and immediately placed into a closable box containing crushed ice. All samples were prepared for the analysis by the working procedures mentioned bellow within 1 hour after sample collection and immediately analysed in triplicate.

#### 2.3.1 Sample preparation using TCA

For the analysis using trichloroacetic acid as a precipitating agent (TCA), 100 μL of water and 100 μL of internal standard working solution were mixed in a 1.8 mL Eppendorf tube with 500 μL of whole blood and then deproteinized by adding 300 μL of TCA-agent (2.0 g of trichloroacetic acid in 8.0 mL of water). Samples were vortexed for ~10s, μlaced into the closable box containing crushed ice for 15 min and then centrifuged at 20,000xg for 10 min. Clarified supernatant was transferred to the brown glass HPLC crimp vial (1.8 mL) for analysis. The spiked samples were prepared as mentioned above and 100 μL of appropriate standard solution was used instead of 100 μL water. In the case of the blank samples, 500 μL of water was used instead of 500 μL of whole blood.

#### 2.3.2 Sample preparation using ZnSO_4_/methanol

Samples with precipitating agent zinc sulfate heptahydrate in methanol were prepared and analysed. Water (100 μL) and internal standard working solution (100 μL) were mixed in a 1.8 mL Eppendorf tube with 300 μL of whole blood and then deproteinized by adding 500 μL of ZnSO_4_-agent (300mM ZnSO_4_.7H_2_O mixed with methanol in a ratio of 3:7 v/v). Samples were vortexed for ~10s, placed into the closable box containing crushed ice for 15 min and then centrifuged at 20,000xg for 10 min. Clarified supernatant was transferred to the brown glass HPLC crimp vial (1.8 mL) for analysis. Again, the spiked sample were prepared as mentioned above and 100 μL of appropriate standard solution was used instead of 100 μL water. In the case of the blank samples, 300 μL of water was used instead of 300 μL of whole blood.

### 2.4 LC-MS/MS analysis

Liquid chromatography was performed by a Thermo Scientific Dionex Ultimate 3000. Quaternary Analytical system was consisted of a solvent rack SRD-3600, binary pump HPG-3400RS, autosampler WPS-3000TRS, column compartment TCC-3000RS, diode array detector DAD-3000RS, heated electrospray HESI II probe, mass detector Velos Pro and a PC with LTQ Tune Plus 2.8 and Xcalibur 4.2.47 software.

The LC-MS/MS system was equipped with a chromatographic column Thermo Acclaim C30 (150 x 2.1 mm, 3 μm) maintained at 15°C. Mobile phase A was prepared by diluting 0.6306 g of ammonium formate with 1 L of water containing 175 μL of formic acid (pH 4.0). Mobile phase B was prepared by diluting 0.6306 g of ammonium formate with 1 L of water containing 1750 μL of formic acid (pH 3.0). Mobile phase C was prepared by diluting 0.0568 g of ammonium formate with 10 mL of mobile phase B and 90 mL of acetonitrile. All mobile phases were prepared daily, filtered through a 0.2 μm nylon filter and degassed before use.

Chromatographic separation was achieved at a flow rate 0.4 mL/min using a gradient of mobile phase A, B and C as follows: (I) 100% mobile phase A from 0 to 5 min, (II) 100% mobile phase B from in 5 min, (III) linear gradient increase to 35% C from 5 to 17 min, (IV) equilibration at starting conditions (100% A) from 17 to 25 min. Injection volume was 10 μl.

Mass spectrometry analysis was completed by multiple reaction monitoring (MRM) or single ion monitoring (SIM) in ESI positive mode using the following tune parameters: capillary voltage = 4 kV, desolvation temperature = 350 °C, sheath gas flow rate = 60 arb., auxiliary gas flow rate = 20 arb., transfer capillary temperature = 350 °C, S-lens RF level = 60 %, Front lens = −9 V, Ion time (MRM) = 200 ms, Ion time (SIM) = 200 ms.

### 2.5 Method validation

Linearity of calibration curves constructed from 6 – 9 concentration levels (each concentration level in triplicate) was evaluated using two acceptability criteria: correlation coefficient (R ≥ 0.9900) and quality coefficient (QC ≥ 5.0 %). Instrument detection limit (IDL) and instrument quantitation limit (IQL) were calculated based on signal-to-noise ratio S/N = 3 and S/N = 10, respectively.

Accuracy, precision and process efficiency were assessed using spiked whole blood samples to achieve a concentration increase of all analytes of 10 and 100 μg/L. All spiked samples were prepared by both sample preparation procedures using TCA and ZnSO_4_/methanol, each in triplicate. Accuracy was determined as the relative error (%) and was expressed as recovery: (measured concentration – nominal concentration/nominal concentration)x 100. The precision was determined as the coefficient of variation (%) and was expressed as repeatability: (standard deviation/mean concentration measured) x 100. Process efficiency (%) was determined: (peak area of an analyte in spiked sample/peak area of the same analyte in standard solution) x 100.

Potential carry-over effects were assessed by monitoring the signal in a blank sample analysed immediately after the analysis of the highest calibration solution (500 μg/L).

## 3. Results and discussion

### 3.1 Optimization of chromatographic and detection conditions

The initial chromatographic and detection conditions were taken over from the Thermo Application Note 294 [21]. The method was optimized to achieve the most stable detector response and the lowest limits of quantitation as possible. At first, ion source parameters were tested to prove that stability of detector response is less than 15 %. Then, fundamental ion optic parameters S-lens RF level (from 0 to 70 %; optimum at 60 %) and Front lens voltage (from −15.0 to −5 V; optimum at −9.0 V) were optimized to ensure the most effective ion transmission from the ion source to the mass analyser. Finally, normalised collision energy (from 0 to 100 %), product ion choice and ion time (from 1 to 500 ms) were chosen to achieve the highest detector response and sufficient number of data points (at least 15 points) across a chromatographic peak. Optimal normalised collision energy needed to achieve optimum fragmentation efficiency ranged from 25 to 50 % for almost all analytes excluding B3, B3-AM and IS-B3. Precursor ions of these compounds do not provide any stable product ion after application of any normalised collision energy – precursor ions either do not fragment at all or fragment totally using this linear ion trap mass analyser. Due to this fact, only SIM mode remained to be chosen in this case. With respect to relatively low scan speed (in comparison with e.g., triple quadrupole or time-of-flight mass analysers), mass detection was divided into 4 different scan windows (2.0-7.0 min, 5.0-11.0 min, 11.0-15.5 min and 15.0-20.0 min) during chromatographic analysis to ensure as long ion time for each analyte or internal standard as possible. All optimal detection conditions for each analyte are listed in Table 1, general detection conditions are mentioned at the end of Chapter 2.4.

**Table 1:**
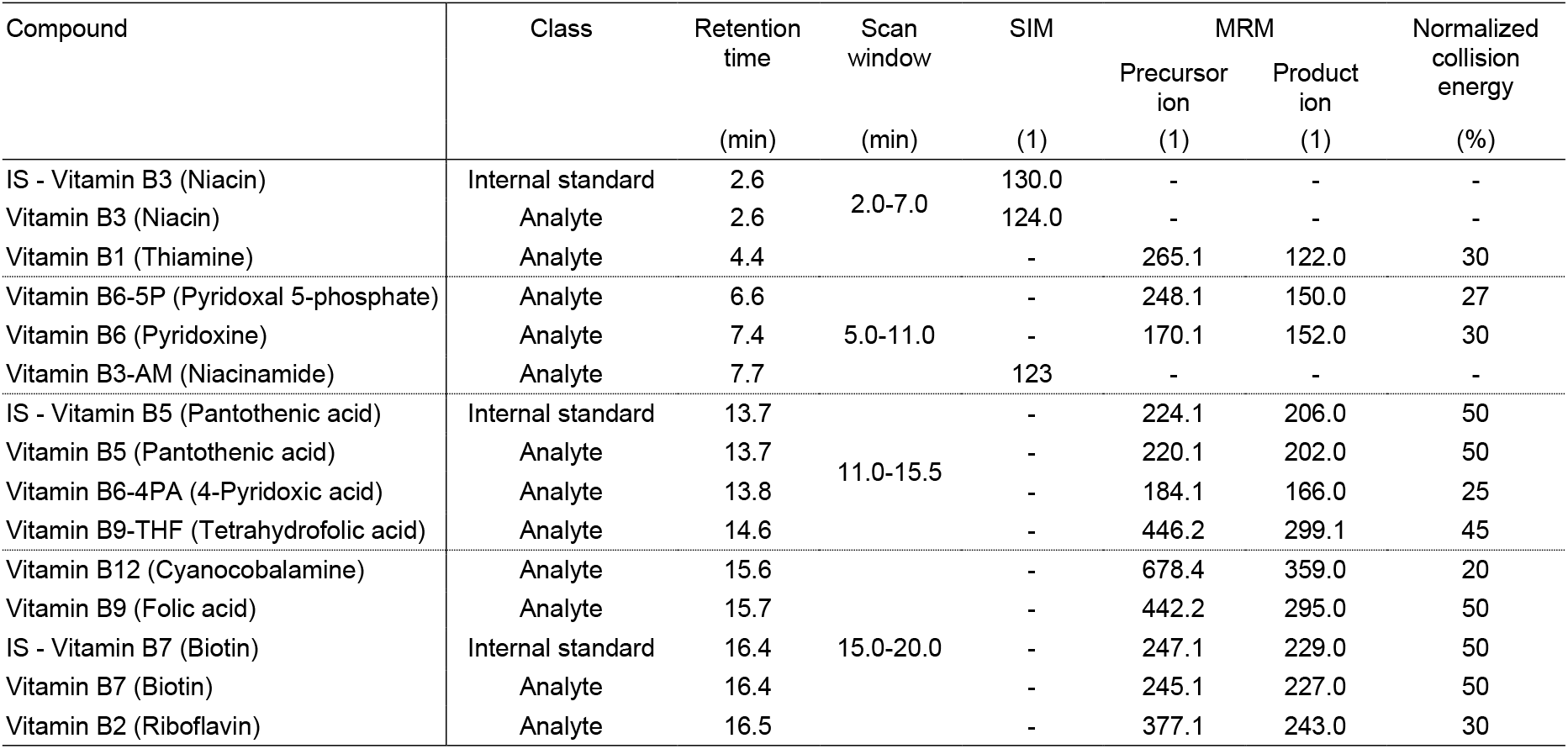
Optimal detection conditions.

### 3.2 Validation characteristics of calibration curves

Calibration curves were measured within the range of 1–500 μg/L, but the whole range of the acceptability criteria (R ≥ 0.9900 and QC ≥ 5.0 %) were fulfilled only for B2, B6, B7 and B12. In the case of other analytes, calibration ranges needed to be narrowed down either from the lowest concentrations (B3, B3-AM, B5 and B6-5P) of from the highest concentrations (B1, B6-4PA or B9). Calibration curve of B9-THF was not assessed due the lack of data caused by insufficient detector response – instrument quantitation limit lies at the highest calibration level (500 μg/L). Correlation coefficients ranged within very good interval of 0.9997-1.0000, quality coefficients ranged from 0.53 to 3.4 %. The best validation characteristics were obtained for B5 (R = 1.0000, QC = 0.53 %), the worst results were achieved for B9 (R = 0.9997, QC = 3.4 %). Instrument quantitation limits ranged from 0.42 to 4.5 μg/L except for B9-THF (500 μg/L). Validation data are listed in the Table 2.

**Table 2:** Validation characteristics of calibration curves.

| Analyte | Internal<br>standard | Instrument<br>detection<br>limit | Instrument<br>quantitation<br>limit | Range | Correlation<br>coefficient | Quality<br>coefficient |
| --- | --- | --- | --- | --- | --- | --- |
|  | (-) | (µg/L) | (µg/L) | (µg/L) | (1) | (1) |
| Vitamin B1 (Thiamine) | IS-B3 | 0.13 | 0.42 | 1 - 100 | 0.9999 | 2.0 |
| Vitamin B2 (Riboflavin) | IS-B7 | 0.27 | 0.91 | 1 - 500 | 1.0000 | 1.3 |
| Vitamin B3 (Niacin) | IS-B3 | 1.2 | 3.8 | 10 - 500 | 0.9999 | 1.7 |
| Vitamin B3-AM (Niacinamide) | IS-B3 | 1.5 | 5.0 | 5 - 500 | 0.9998 | 2.7 |
| Vitamin B5 (Pantothenic acid) | IS-B5 | 0.18 | 0.56 | 2.5 - 500 | 1.0000 | 0.53 |
| Vitamin B6 (Pyridoxine) | IS-B5 | 0.14 | 0.45 | 1 - 500 | 0.9998 | 2.9 |
| Vitamin B6-5P (Pyridoxal 5-phosphate) | IS-B3 | 1.4 | 4.5 | 5 - 500 | 0.9999 | 1.8 |
| Vitamin B6-4PA (4-Pyridoxic acid) | IS-B5 | 0.30 | 1.0 | 1 - 100 | 0.9997 | 3.2 |
| Vitamin B7 (Biotin) | IS-B7 | 0.27 | 0.91 | 1 - 500 | 1.0000 | 1.3 |
| Vitamin B9 (Folic acid) | IS-B7 | 0.21 | 0.71 | 1 - 100 | 0.9997 | 3.4 |
| Vitamin B9-THF (Tetrahydrofolic acid) | IS-B7 | 150 | 500 | n.a. | n.a. | n.a. |
| Vitamin B12 (Cyanocobalamine) | IS-B5 | 0.30 | 1.0 | 1 - 500 | 0.9999 | 2.2 |

### 3.3 Effect of protein-precipitating agents on accuracy, precision and process efficiency of the method

Protein-precipitating agent could play a crucial role in sample pre-treatment for trace analysis. Whole blood is one of the most challenging matrixes for LC/MS technique due to negative ionization effects during electrospray ionization. Results of comprehensive study [22] focused on optimization of protein precipitation from plasma samples based upon effectiveness of protein removal and ionization effect in LC-MS/MS shows that 10 % TCA is an optimal protein-precipitating agent for removal of plasma proteins, together with minimal ionization effect. Nevertheless, whole blood is more complex matrix than plasma and sample pre-treatment step must overcome not only sufficient protein precipitation but also haemolysis step for an efficient release of analytes presented in erythrocytes. Finally, physicochemical properties (e.g., pH) of a protein-precipitating agent have to be taken into consideration with regard to the analytes stability. One of the very effective non-acidic agents for whole blood precipitation is 0.2 M zinc sulphate solution in organic solvent [23, 24]. Both sample preparation methods mentioned above were assessed for presented determination of B vitamins in whole blood using LC/MS in this article.

Protein precipitation using ZnSO_4_/methanol provides better results at both spike levels 10 and 100 μg/L in general, but B6-5P, B9 and B9-THF were not detected regardless of used protein-precipitating agent. Ranges of evaluated parameters (accuracy, precision and process efficiency) of the remaining analytes were narrower and closer to optimal values in almost all cases except for precision at 100 μg/L (1.0-13 % for ZnSO_4_/methanol, but 1.6-6.8 % for TCA). Worse precision in case of protein precipitation using ZnSO_4_/methanol was observed for B1 at both spike levels – 9.9 % at 10 μg/L (TCA 4.0 %) and 13 % at 100 μg/L (TCA 1.6 %). This higher variableness of precision values is caused by peak broadening due to the high percentage of organic solvent in ZnSO_4_/methanol solution (70 %), relatively high injection volume (10 μL) and narrow chromatographic column (2.1 mm). Excluding B1 results, precision ranged from 0.50 to 7.1 % at 10 μg/L and from 1.0 to 6.2 at 10 μg/L.

The choice of the precipitating agent also significantly affects the accuracy ot the method. In the case of TCA agent the accuracy ranged from 26 to 122 % at 10 μg/L and from 30 to 112 % at 100 μg/L but better results were obtained in the case of ZnSO_4_/methanol – from 90 to 114 % at 10 μg/L and from 89 to 120 % at 100 μg/L. Low accuracy when using TCA is noticeable for three B vitamins: B12 (26 % at 10 μg/L and 30 % at 100 μg/L), B2 (51 % at 10 μg/L and 56 % at 100 μg/L) and B5 (55 % at 10 μg/L and 68 % at 100 μg/L).

Process efficiency (PE) assesses LC-MS method as a whole because this parameter takes into consideration not only contributions of matrix effects but also efficiency (recovery) of the extraction process. The method using TCA agent for protein precipitation step shows rather big differences at both spike levels 10 and 100 μg/L in this parameter – PE of three B vitamins were higher than 150 % (B1, B3 and B6), PE of three B vitamins were lower than 50 % (B2, B7 and B12) and only three B vitamins provides PE between 50-150 % (B3-AM: 94 and 101 %; B5: 111 and 117 %; B6-4PA: 100 and 107 %). Much better results were obtained using the ZnSO_4_/methanol method because PE were ranging from 65 to 108 % at both spike levels 10 and 100 μg/L for all nine B vitamins (B1, B2, B3, B3-AM, B5, B6, B6-4PA, B7 and B12).

### 3.4 Stability of B vitamins in whole blood samples

The stability of B vitamins in the whole blood samples spiked at 100 μg/L level was evaluated at −18°C to check whether a short-time storage of blood in collection tubes in an ordinary freezer does not affect concentration of analytes. The stability of B6 is decreased at such conditions, 17 % content decrease was observed after one day, and 51 % content decrease in four-day storage time. The content of B1, B2, B3, B5, B6-PA, B7 and B12 was stable for the first three days, on the fourth day a decrease of 5-23 % was observed. Such results correspond to the previously reported observations of stability of B vitamins in whole blood [25], B1 in the erythrocytes [20] or vitamins B9 and B12 in human serum [26]. Slight differences may be due to EDTA presence in the previous studies. The content of B3-AM could not be evaluated due to the high detector response that lies in the range of calibration curve due to its extremely high concentration in the tested whole blood sample. However, B3-AM, as well as B3, are generally known for a sufficient stability at various conditions and their activity is not affected by heat, light, acid, alkali or oxidation [27]. Remaining B vitamins (B6-5P, B9 and B9-THF) were not detected due to unsuccessful validation characteristics of the tested sample preparation methods using trichloroacetic acid (Table 3) or ZnSO_4_/methanol (Table 4) as a precipitating agent.

**Table 3:** Validation characteristics of the method using TCA as a precipitating agent.

| Analyte | Spike 10 µg/L |  |  | Spike 100 µg/L |  |  |
| --- | --- | --- | --- | --- | --- | --- |
|  | Accuracy | Precision | Process efficiency | Accuracy | Precision | Process efficiency |
|  | (%) | (%) | (%) | (%) | (%) | (%) |
| Vitamin B1 (Thiamine) | 73 | 4.0 | 229 | 78 | 1.6 | 166 |
| Vitamin B2 (Riboflavin) | 51 | 1.8 | 29 | 56 | 2.3 | 30 |
| Vitamin B3 (Niacin) | 91 | 8.4 | 250 | 102 | 5.7 | 199 |
| Vitamin B3-AM (Niacinamide) | 71 | 6.2 | 94 | 79 | 4.8 | 101 |
| Vitamin B5 (Pantothenic acid) | 55 | 2.7 | 117 | 68 | 1.6 | 111 |
| Vitamin B6 (Pyridoxine) | 122 | 7.1 | 156 | 112 | 4.7 | 151 |
| Vitamin B6-5P (Pyridoxal 5-phosphate) | < 1 | n.a. | n.a. | < 1 | n.a. | n.a. |
| Vitamin B6-4PA (4-Pyridoxic acid) | 88 | 5.8 | 100 | 85 | 2.2 | 107 |
| Vitamin B7 (Biotin) | 93 | 2.9 | 44 | 95 | 2.5 | 61 |
| Vitamin B9 (Folic acid) | < 1 | n.a. | n.a. | < 1 | n.a. | n.a. |
| Vitamin B9-THF (Tetrahydrofolic acid) | < 1 | n.a. | n.a. | < 1 | n.a. | n.a. |
| Vitamin B12 (Cyanocobalamine) | 26 | 24 | 41 | 30 | 6.8 | 43 |
| Range | 26–122 | 1,8–24 | 29–250 | 30–112 | 1.6–6.8 | 30–199 |

**Table 4:** Validation characteristics of the method using ZnSO_4_/methanol as a precipitating agent.

| Analyte | Spike 10 µg/L |  |  | Spike 100 µg/L |  |  |
| --- | --- | --- | --- | --- | --- | --- |
|  | Accuracy | Precision | Process efficiency | Accuracy | Precision | Process efficiency |
|  | (%) | (%) | (%) | (%) | (%) | (%) |
| Vitamin B1 (Thiamine) | 99 | 9.9 | 96 | 116 | 13 | 75 |
| Vitamin B2 (Riboflavin) | 108 | 3.4 | 87 | 105 | 3.6 | 75 |
| Vitamin B3 (Niacin) | 100 | 7.0 | 91 | 89 | 6.2 | 65 |
| Vitamin B3-AM (Niacinamide) | 92 | 5.3 | 82 | 90 | 5.5 | 96 |
| Vitamin B5 (Pantothenic acid) | 91 | 1.2 | 86 | 92 | 1.4 | 81 |
| Vitamin B6 (Pyridoxine) | 114 | 0.50 | 84 | 120 | 2.5 | 100 |
| Vitamin B6-5P (Pyridoxal 5-phosphate) | < 1 | n.a. | n.a. | < 1 | n.a. | n.a. |
| Vitamin B6-4PA (4-Pyridoxic acid) | 105 | 1.7 | 90 | 104 | 1.0 | 91 |
| Vitamin B7 (Biotin) | 90 | 3.8 | 71 | 96 | 2.6 | 69 |
| Vitamin B9 (Folic acid) | < 1 | n.a. | n.a. | < 1 | n.a. | n.a. |
| Vitamin B9-THF (Tetrahydrofolic acid) | < 1 | n.a. | n.a. | < 1 | n.a. | n.a. |
| Vitamin B12 (Cyanocobalamine) | 99 | 7.1 | 97 | 112 | 3.7 | 108 |
| Range | 90-114 | 0.50-9.9 | 71–97 | 89–120 | 1.0–13 | 65–108 |

## 4. Conclusions

The aim of this work was to quantify simultaneously seven B vitamins and two B vitamin derivatives in a whole blood. The suitability of two different protein precipitating agents was compared using ZnSO_4_/methanol and trichloroacetic acid. Attention was focused on detailed optimization of mass detection using linear ion trap mass analyser for rapid and robust LC-MS/MS method. The usage of ZnSO_4_/methanol instead of trichloroacetic acid for protein precipitation step provides high percentages of accuracy, precision and process efficiency. As far as we know, this is the first method that allows to quantify thiamine, riboflavin, niacin, niacinamide, pantothenic acid, pyridoxine, 4-pyridoxic acid, biotin and cyanocobalamine in a whole blood simultaneously.

**Figure 1:**
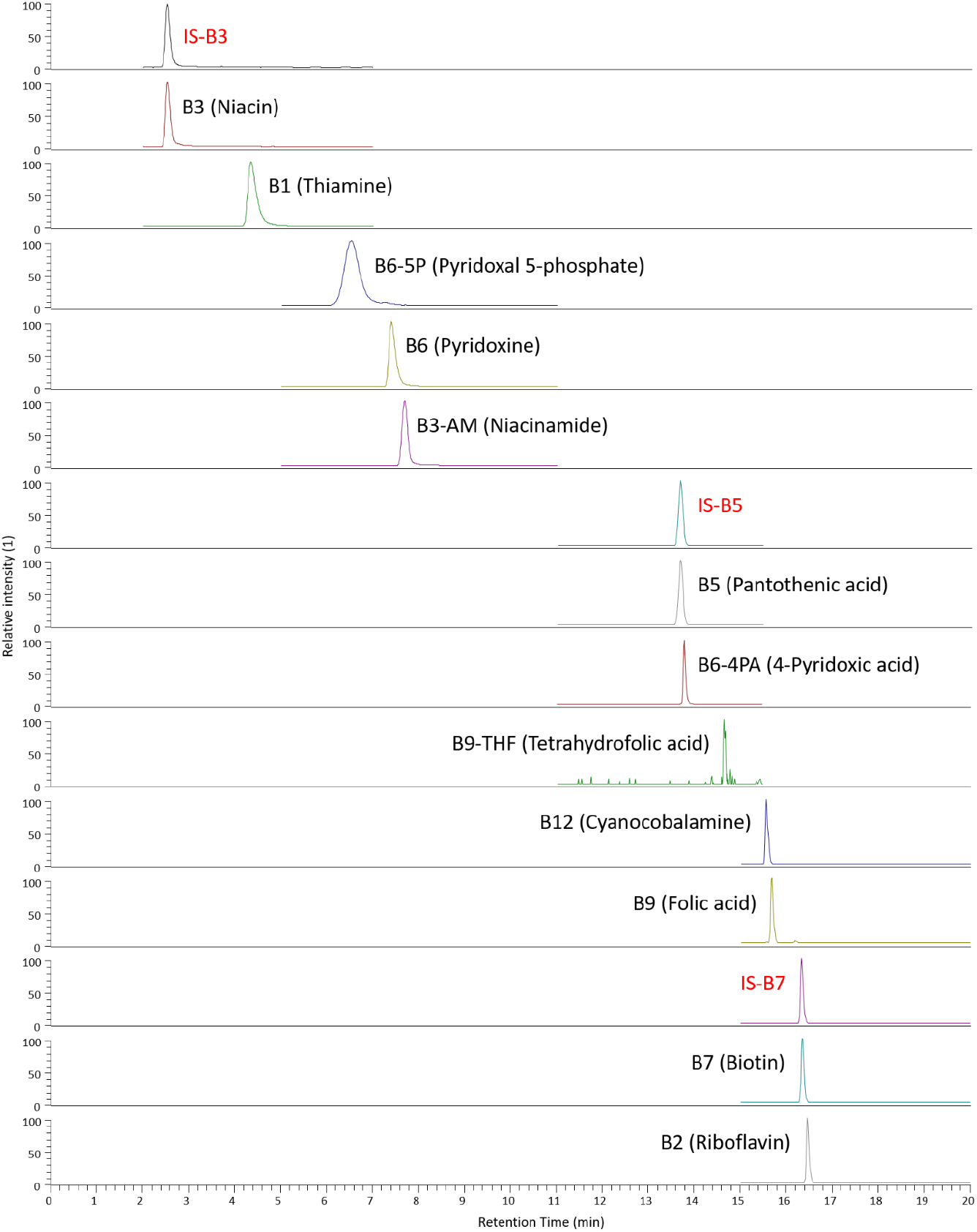
Representative chromatogram of the highest calibration solution (500 μg/L).

## Declaration of competing interest

All authors declare no conflict of interest.

## Acknowledgements

The authors gratefully acknowledge the financial support of the research provided by the grant project No. 18-07711S sponsored by the Czech Science Foundation.

## Notes

### Competing Interest Statement

The authors have declared no competing interest.

